# Protein-ligand interfaces are polarized: Discovery of a strong trend for intermolecular hydrogen bonds to favor donors on the protein side with implications for predicting and designing ligand complexes

**DOI:** 10.1101/260612

**Authors:** Sebastian Raschka, Alex J. Wolf, Joseph Bemister-Buffington, Leslie A. Kuhn

## Abstract

Understanding how proteins encode ligand specificity is fascinating and similar in importance to deciphering the genetic code. For protein-ligand recognition, the combination of an almost infinite variety of interfacial shapes and patterns of chemical groups makes the problem especially challenging. Here we analyze data across non-homologous proteins in complex with small biological ligands to address observations made in our inhibitor discovery projects: that proteins favor donating H-bonds to ligands and avoid using groups with both H-bond donor and acceptor capacity. The resulting clear and significant chemical group matching preferences elucidate the code for protein-native ligand binding, similar to the dominant patterns found in nucleic acid base-pairing. On average, 90% of the keto and carboxylate oxygens occurring in the biological ligands formed direct H-bonds to the protein. A two-fold preference was found for protein atoms to act as H-bond donors and ligand atoms to act as acceptors, and 76% of all intermolecular H-bonds involved an amine donor. Together, the tight chemical and geometric constraints associated with satisfying donor groups generate a hydrogen-bonding lock that can be matched only by ligands bearing the right acceptor-rich key. Measuring an index of H-bond preference based on the observed chemical trends proved sufficient to predict other protein-ligand complexes and can be used to guide molecular design. The resulting Hbind and Protein Recognition Index software packages are being made available for rigorously defining intermolecular H-bonds and measuring the extent to which H-bonding patterns in a given complex match the preference key.

**Abbreviations:**

3D: three-dimensional
CATH: Class Architecture Topology Homologous superfamily
H-bonds: hydrogen bonds
MMFF94: Merck Molecular Force Field
PDB: Protein Data Bank
PRI: Protein Recognition Index

## Introduction

Across several molecular docking, alignment, screening and crystallographic data analysis projects [1–4], we made the following observations:

- Molecules enhanced in chemical groups having both hydrogen bond (H-bond) donor and acceptor capacity (e.g., hydroxyl groups) tend to lead to false-positive rankings in molecular screening and inaccurate prediction of binding poses for known ligands. This is apparently due to the greater number of potential favorable interactions of donor + acceptor matches (which are augmented by the bond-rotational possibilities for hydroxyl groups), leading to higher protein-ligand interaction counts and overestimated affinity, relative to ligands enhanced in donor-only and acceptor-only groups.
- When analyzing the protein binding sites in a number of protein-small molecule crystal structures, we also noticed that H-bonds tended to be donated from the protein to the ligand, rather than observing an even distribution of donors and acceptors on both sides of the interface.
- While optimizing the docking scoring function for SLIDE [1] and the surface alignment scoring function for ArtSurf [4] by training on known complexes or site matches, we noted that the terms for matching chemical groups with both donor and acceptor capacity received much smaller weights than the weights for matching donor-only or acceptor-only groups.

An interesting possibility is that nature avoids the presence of chemical groups bearing both H-bond donor and acceptor capacity, such as hydroxyl groups, in the binding sites of proteins or ligands. The many ways of satisfying these groups with H-bond partners could lead to non-selective ligand binding. This hypothesis appears to be supported by the second observation that proteins selectively donate (rather than donate and accept) H-bonds to small molecules. Since those observations were made anecdotally over time and may not hold for protein-ligand complexes in general, the present study was designed to assess whether the above trends (or others) are consistently present in a set of 136 non-homologous proteins bound to a range of biologically relevant small molecules. Selecting this set of proteins with no binding site structural homology between any constituent pair removed potential bias towards a given fold, sequence, or function. We then tested whether the resulting statistics of H-bonding trends alone provided enough information to predict the orientation of ligands relative to their protein partners. The goal was to evaluate whether trends derived from many complexes hold for individual examples well enough to predict the native interactions. Predicting the orientation of ligands for 30 additional complexes also addressed whether the observed trends constitute an essential part of the code for recognition between proteins and ligands.

Advances over the decades in our understanding of protein H-bonding have been well-reviewed in [5]. The literature most relevant to the present study falls into two areas: defining energetically favorable H-bonds in terms of geometry (given the integral relationship between favorable geometry and favorable energy) and characterizing H-bond interactions in protein-ligand complexes. Nittinger et al. [5] analyzed a large number of protein-ligand structures to define preferred H-bond geometries and the extent to which H-bonds observed in experimental structures match theoretically predicted H-bonds based on the valence shell electron pair repulsion model. Their focus was on furthering the accurate modeling and parameterization of H-bonds. As in the work of McDonald & Thornton [6], Nittinger et al. found only small energetic differences in out-of-plane H-bonding angles for sp^2^ groups such as keto oxygens. This has a key impact on ligand orientational selectivity for donor versus acceptor groups in the present work. Panigrahi and Desiraju [7] also studied protein-ligand H-bonds across a number of diverse, if not necessarily non-homologous, small molecule complexes. Their criteria for defining H-bonds (proton within 3.0 Å of acceptor, resulting in a donor-acceptor distance of up to 4 Å, and donor-H-acceptor angle greater than 90˚, with 90˚ reflecting a very weak H-bond) were less stringent than those used here, which could result in the inclusion of relatively low-strength, second-shell (less direct) interactions in their statistics. They defined strong H-bonds as those involving polar donor and acceptor atoms, versus weak H-bonds formed by CH donors to oxygen acceptors. They found that N-H---O and O-H---O H-bonds tended to be linear, C-H---O H-bonds to oxygen with Gly and Tyr as donors were ubiquitous in active sites, and that ligands accept twice as often as they donate H-bonds to the protein, consistent with Lipinski’s *Rule of 5* [8]. The current work focuses on identifying chemical interaction patterns between proteins and their ligands at an atomic chemistry rather than functional group scale, evaluating underlying reasons for such patterns, including ligand selectivity, and testing the extent to which these patterns can predict native interactions.

## Methods

### Dataset

A dataset of well-resolved protein complexes with biologically relevant small molecules was constructed based on the intersection between proteins representing different CATH structural folds [9] (<u>C</u>lass, <u>A</u>rchitecture, <u>T</u>opology, <u>H</u>omologous superfamily; http://www.cathdb.info) and a set of well-resolved protein structures bound to small organic molecules with known affinity from Binding MOAD [10] (http://bindingmoad.org). This resulted in a dataset of 136 non-homologous protein structures (Supplementary Material 1, Table S1) from the Protein Data Bank [11] (PDB; http://www.rcsb.org) with a resolution of 2.4 Å or better; 90% of the structures were solved at 2.0 Å resolution or better. The protein structures were bound to a diverse set of small ligands (25 peptides, 50 nucleotides, bases and base analogs, and 61 other organic molecules). None of the structures were problematic in ligand fitting or resolution according to the Iridium quality analysis of protein-ligand fitting and refinement [12].

### Protonation

Protonation of each protein-ligand complex was performed with the OptHyd method in YASARA Structure [13] (version 16.4.6; http://www.yasara.org; see details in Supplementary Material 2, supp_material_2_yasara_opthyd_macro.txt), retaining interfacial metals and removing bound water molecules, with the goal of assessing direct, strong interactions between proteins and their ligands. During the addition of hydrogen atoms and optimization of the H-bond network using YASARA, heavy atom positions were maintained except for the rotation of the terminal amide groups of asparagine and glutamine side chains through 180˚ when interchange of the =O and −NH_2_ groups resulted in improvement in polar interactions. This step disambiguates the fitting of these side chains into electron density due to the similar density of oxygen and nitrogen atoms at typical crystallographic resolution. YASARA assigns the tautomeric state of the imidazole groups in histidine side chains according to the intra- and intermolecular hydrogen and metal bonding of each histidine and the influence of neighboring polar groups on the pK_a_ of the imidazole ring [14]. For nucleotidyl ligands and bases, the results of high-level *ab initio* calculations on protonation states and tautomers, and how they are influenced by H-bonding in complexes, were also considered [15].

### Optimization of proton orientation

To minimize steric clashes and optimize the polar interaction network, OptHyd also optimized the orientation of protein and ligand protons (for instance, the hydrogen positions in rotatable NH_3_ and OH groups). This method uses the YAMBER force field, a second-generation force field derived from AMBER, which was self-parameterized according to the protonated protein, water molecules, and ions present in the complete unit cells of 50 high resolution X-ray structures [16]. All 136 of the complexes in our analysis were checked for agreement between YASARA protonation of the ligand in complex with the protein, relative to protonation of the ligand alone using molcharge in OpenEye QUACPAC with the AM1-BCC option [17] (version 1.7.0.2; https://www.eyesopen.com/quacpac; OpenEye Scientific Software, Santa Fe, NM). Protonation and ligand valences resulting from YASARA were also visually inspected with PyMOL [18] (version 1.8.2.2, https://www.schrodinger.com/pymol). In cases of ambiguities or differences in protonation state or valence, the chemical literature for the protein-ligand complex and protonation studies for the ligand were consulted, resulting in manual correction relative to the YASARA protonation in a few cases. The protonated ligands provided in PDB format by YASARA were converted to Tripos MOL2 format with the OpenEye OEChem toolkit (version 1.7.2.4; https://www.eyesopen.com/oechem-tk; OpenEye Scientific Software, Santa Fe, NM).

### Influence of partial charges

In the Hbind software developed in this work to define H-bond interactions (including salt bridges satisfying H-bond criteria, described in the following paragraph), partial charges are only used to assess whether an atom pair can form longer-range salt bridges. The salt bridge assignment requires a higher than 0.3 charge magnitude on the ligand atom interacting with a charged protein or metal atom and a maximum distance of 4.5 Å between the donor and acceptor. These longer-range salt bridges are often second-shell interactions and thus were not included in the current analysis. Hence, as expected, charges assigned by either the Merck Molecular Force Field [19] (MMFF94) or AM1-BCC [17] using molcharge in QUACPAC resulted in the same list of direct H-bond and metal bridge interactions for the 136 complexes. The protonated ligands with MMFF94 charges are provided (in the same reference frame as the corresponding PDB complex) in a multi-MOL2 file [20] in Supplementary Material 3 (supp_material_3_lig_in_orig_PDB_refframe.mol2).

### Hbind software

This software developed in our laboratory (available from GitHub at https://github.com/psa-lab/Hbind) was used to define direct H-bonds and metal bonds with ligands. Pauling wrote, “Only the most electronegative atoms should form H-bonds, and the strength of the bond should increase with increase in the electronegativity of the two bonded atoms…[Thus] we might expect that fluorine, oxygen, nitrogen and chlorine would possess this ability, to an extent decreasing in this order.” [21]. In our software, nitrogen and oxygen atoms are considered as potential donors or acceptors of H-bonds and fluorine and chlorine as potential acceptors. Hbind interprets the donor/acceptor capacity of ligand atoms from information in the MOL2 file detailing the hybridization, the order of covalent bonds with neighboring atoms, and the protonation state of these atoms. The software implicitly evaluates by analytic geometry all orientations of protons in rotatable groups for their ability to satisfy the H-bond criteria defined below, while not altering their coordinates in the PDB or MOL2 file. For instance, protons in XNH3 and X-OH groups can adopt any sterically admissible position on a circle upon rotation of the X-N or X-O single bond. The H-bond identification criteria are based on those of Ippolito et al. [22] and McDonald and Thornton [6], all of which must be met:

- Hydrogen to acceptor distance: 1.5-2.5 Å
- Donor to acceptor distance: 2.4-3.5 Å
- Donor-H-acceptor angle (Θ): 120-180**°**
- Pre-acceptor–acceptor–H angle (ϕ): 90-180°

These donor, hydrogen, and acceptor geometries are depicted in Fig. 1.

**Fig. 1.**
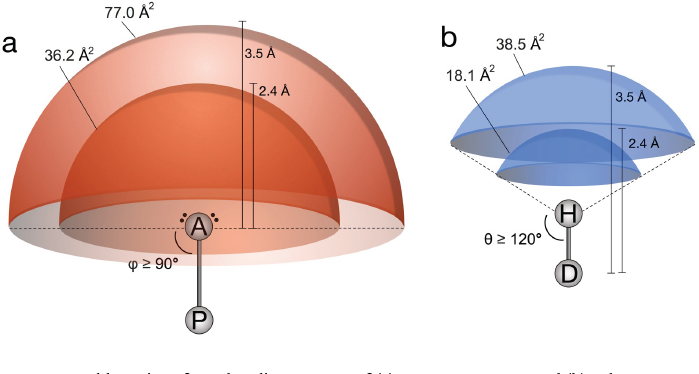
Favorable regions for H-bonding partners of (a) an acceptor atom, and (b) a donor atom in the protein or ligand. Based on the geometric criteria described in the text, the outer and inner shells represent the maximum and minimum distances for the H-bond partner atom relative to the acceptor (a) or donor (b). Given that the pre-acceptor–acceptor–H angle (ϕ) can range from 90-180˚, the surface area of the outer shell defining the maximum distance at which a donor atom can interact favorably, within 3.5 Å of the acceptor, is 77.0 Å^2^. The inner shell at 2.4 Å correspondingly represents the surface of minimum distance for the donor relative to the acceptor. The favorable volume for a protein atom to H-bond with a donor atom (a) or an acceptor atom (b) on the ligand is defined as the volume between the inner and outer shells. Because the pre-acceptor–acceptor–H angle (ϕ) can range from 90-180˚ for each lone pair on the acceptor (a), while the range for the donor-H-acceptor angle in (b) is narrower (120-180˚), the volume in which a ligand proton can favorably bind to a protein acceptor atom (60.6 Å^3^) is twice the volume where an acceptor atom can make a favorable interaction with a donor atom (30.3 Å^3^).

The following criteria were used for protein or ligand-bound metals to form a bond with an atom on the second molecule bearing a lone pair of electrons:

- Lone pair atom distance to K or Na: 2.0-2.9 Å
- Lone pair atom distance to Ca, Co, Cu, Fe, Mg, Mn, Ni, or Zn: 1.7-2.6 Å

Hbind calculates and outputs the interaction distance and angles between each protein-ligand atom pair forming an H-bond or metal interaction. Additional command-line options are available to list longer-range salt bridges (up to 4.5 Å between protein and ligand), direct hydrophobic contacts, and the protein-ligand orientation and affinity scores and terms calculated by SLIDE [1] (version 3.4, http://kuhnlab.bmb.msu.edu/software/slide/index.html).

### Identification of ligand H-bonding patterns

This analysis aimed to identify any consistent patterns of nitrogen donor interactions from proteins to ligands in the dataset of 136 non-homologous complexes. When visualizing the complexes with PyMOL, geometrical similarities were apparent in the H-bond networks with nucleotidyl ligands, involving a visually distinctive pattern of protein H-bond donors. To assess this objectively, the pattern of H-bond interactions within each complex was represented by a binary vector listing the presence (1) or absence (0) of an H-bond to the ligand for each position in the sequence. Because the number of possible interaction patterns for protein sequences with hundreds of residues and arbitrary spacing between the H-bonding positions is almost infinite, we chose to focus on the subcase of identifying local H-bonding sequence patterns with at least three interacting residues and no more than 10 residues intervening between a pair of successive interactions. For each protein, the initial H-bonding vector was then split into non-overlapping sub-vectors (local motifs), such that each sub-vector started and ended with an H-bonding residue and did not contain a contiguous subsequence of more than ten zero-elements (non-interacting residues). For example, an H-bond interaction vector consisting only of nitrogen donors (here, from PDB entry 1f5n [23]) appears as follows between the vertical bars, where the initial number (1) is the first residue number in the sequence, and the last number (1361) is the final residue number:

1| 0 0 0… 1 1 1 1 1 0 0 0 0 0 0 0 0 0 0 0 0 0 0 1 0 0 0 0 0 1 1… 0 0 0 |1361

Note that these vectors are formatted as binary sequences, where residues forming interfacial H-bonds are labeled with 1’s, and residues not involved in H-bonds are set to 0. The following excerpt of a protein H-bond interaction vector shows a region within the above complete vector containing several local interaction vectors:

43| 1 1 1 1 1 0 0 0 0 0 0 0 0 0 0 0 0 0 0 1 0 0 0 0 0 1 1|69

The extracted sub-vectors or local H-bonding motifs were then:

43| 1 1 1 1 1 |47 *and* 62| 1 0 0 0 0 0 1 1 |69

Once all sub-vectors were extracted, they were tabulated by protein and concatenated into a dataset containing the local motifs from all 136 complexes. Trailing zeros were added to facilitate displaying the results:

N_1B5E_1_D400| 1 0 1 1 0 0 0 0 0 0 0 0 0 0 0 0

N_1BX4_1_A350| 1 1 0 0 1 0 0 0 0 0 0 0 0 0 0 0

N_1CIP_1_A355| 1 1 1 1 0 0 0 0 0 0 0 0 0 0 0 0

N_1F0L_1_A601| 1 1 0 1 0 0 0 0 0 0 0 0 0 0 0 0

N_1F5N_1_A593| 1 1 1 1 1 0 0 0 0 0 0 0 0 0 0 0

N_1F5N_2_A593| 1 0 0 0 0 0 1 1 0 0 0 0 0 0 0 0

*etc.*

The first letter in each row denotes whether this subsequence corresponds to a peptide-like (P), nucleotide-like (N), or other organic (O) ligand. Because some protein complexes contained more than one interaction motif, the digit following the underscore after the PDB code indexes the motifs in a given protein. The first character after the next underscore is the PDB chain ID of the protein and ligand analyzed, and the remaining digits specify the residue number of the ligand molecule.

### Software for statistical analyses

The parsing of Hbind interaction tables and the statistical analyses in this work were carried out in Python using NumPy [24] (version 1.13.3, http://www.numpy.org), SciPy [25] (version 0.19.1, https://www.scipy.org), and Pandas [26] (version 0.20.3, https://pandas.pydata.org). The BioPandas package [27] (version 0.2.2, http://rasbt.github.io/biopandas/) was used to compute statistics from MOL2 and PDB files.

### Visualization and plotting software

All data plots were created using the matplotlib library [28] (version 2.0.2, https://matplotlib.org). The Affinity Designer software (version 1.6.0, https://affinity.serif.com/en-us/designer/) was used to enhance the readability of figure labels, as necessary. Structural renderings of molecules were created in PyMOL (version 1.8.2.2, https://pymol.org), and figures depicting geometric properties were drawn in OmniGraffle (version 7.5, https://www.omnigroup.com/omnigraffle).

## Results and Discussion

Before data analysis, the valences and protonation state were carefully checked for each of the 136 ligand complexes relative to the chemical literature. As described in the Methods, the free ligand structures were protonated with OpenEye molcharge and the protein-bound ligand structures were protonated separately with the YASARA OptHyd protocol, since the quantum chemistry calculations in molcharge would not be feasible for entire protein-ligand complexes. However, the resulting differences in protonation between the apo and bound ligand states were dominated by the ability of the respective software to correctly process the ligand structure from the PDB file (inferring atom hybridization, valence, and the charge state of polar atoms) rather than reflecting differences in protonation due to the ligand being in a protein-bound or free state. To address the interesting question of how often the protonation state is the same in the apo versus protein-bound ligand, the YASARA OptHyd protocol was also run on each ligand in the free state, and its proton assignment was compared with the YASARA results on the 136 complexes. The results were:

- 56 of the 136 cases (41.2%) had the same protonation and essentially the same proton orientation when the ligand was protonated as a separate molecule or in complex with the protein
- 52 of the 136 cases (38.2%) had the same protonation with one or more protons in a different orientation
- 18 of the 136 cases (13.2%) had a different protonation state between the apo and bound forms of the ligand; this was often due to mis-protonation of a phosphate group in the apo state or the presence of a metal ion in the ligand binding site, requiring deprotonation of the ligand atom in order to ligate the metal
- 7 of the 136 (5.1%) had a flipped amide or imidazole group between the apo and bound states to optimize hydrogen bonding. In one case, such a flip would be biologically unlikely due to electron delocalization over the bond connecting the amide to an aromatic ring; the resulting partial double bond character presents a high energetic barrier to flipping
- 3 of the 136 cases (2.2%) involved protonation differences due to the ligand bond structure being handled incorrectly

In 8 of the 136 complexes that were analyzed in the following sections, a proton needed to be added to the ligand, post-YASARA OptHyd protonation of the protein-ligand complex, often due to incomplete specification of the ligand bond structure in the PDB file interpreted by YASARA. In 11 of the 136 cases, a proton needed to be deleted from the ligand for the same reason. A single ligand containing boron also required manual correction. Though 79% (free state) or 86% (bound state) of the ligands were protonated correctly by the automated YASARA OptHyd procedure, these results highlight the importance of manually checking the correctness of the structure and protonation state of ligands before further analysis.

The output of Hbind with direct intermolecular H-bonds and metal interactions for all 136 correctly protonated complexes (Supplementary Material 1, Table S1) was the basis for addressing a series of molecular recognition questions presented and discussed in this section. The complete Hbind interaction data is provided in Supplementary Material file 4 (supp_material_4_hbind_interaction_tables.txt), with an example for one complex shown in Fig. 2. We then addressed the following questions, to quantify and understand patterns in interfacial hydrogen bonding between proteins and their small biological ligands.

**Fig. 2.**
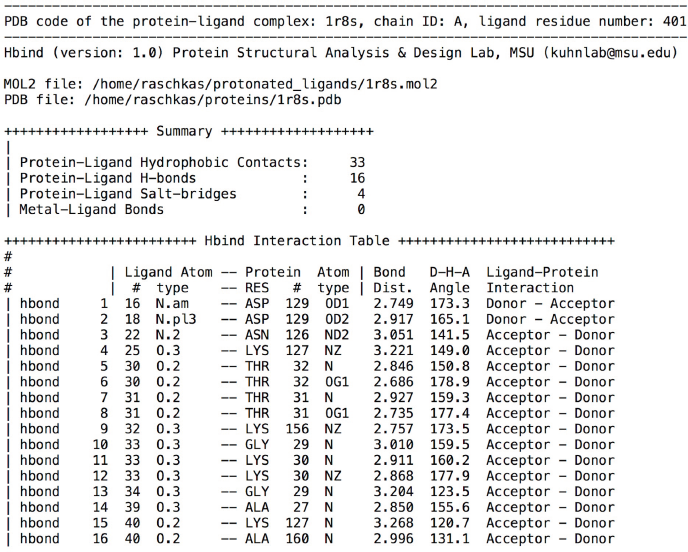
Example of Hbind intermolecular direct H-bond and metal interaction output for chain A of PDB entry 1r8s in complex with ligand GDP (chain ID: A, ligand residue number: 401), showing only those interactions meeting the criteria defined in the Methods. The ligand atom number and type are from the MOL2 file definition, and the protein residue number and atom type, bond length between H-bond donor and acceptor atoms, and the donor-hydrogen-acceptor (θ) angle are also listed. The final columns indicate the orientation of the hydrogen bond, i.e., whether the ligand or protein contributed the donor atom, and likewise for the acceptor.

### 1. Are donor groups on proteins preferred for H-bonding to biological ligands?

The interaction tables for 136 complexes were analyzed to count the frequency of protein atoms acting as H-bond acceptors versus donors in direct H-bonds to the ligand and likewise for ligand atoms (Fig. 3). The preference for the protein to donate H-bonds to a ligand acceptor atom was more than 2:1, with 712 H-bonds donated by the protein to the ligand and 345 H-bonds from the ligand accepted by the protein, across all 136 complexes. Since H-bonds were analyzed based on atomic interactions, including proton positions, a residue or atom could participate in multiple H-bonds if all the angular and distance criteria were met for each bond.

**Fig. 3.**
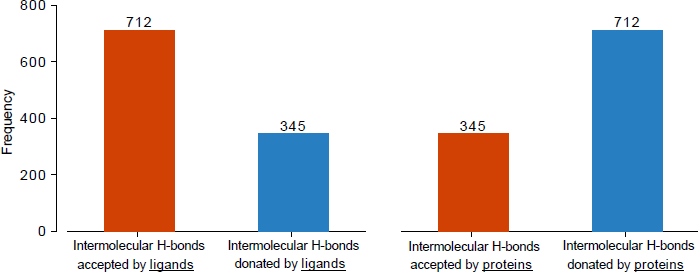
Frequency of donated and accepted intermolecular H-bonds across the 136 diverse complexes shown from the ligand’s perspective (two bars on the left) and the protein’s perspective (two bars on the right). Throughout the figures, red is used to indicate H-bond acceptors, while blue indicates donors.

When subdivided further according to the patterns of nitrogen and oxygen atoms involved in protein-ligand H-bonds, an interesting trend came to light: the majority (70%) involved both a nitrogen atom donor and an oxygen acceptor (Table 1), with a full 76% of intermolecular H-bonds donated by a nitrogen atom. The second most prevalent case paired a hydroxyl group donor with an oxygen acceptor (24%). Other possibilities for native ligand H-bonding were rare, particularly nitrogen atoms acting as H-bond acceptors, whether on the protein or ligand side. The tendency of hydroxyl groups to contribute only one-quarter of all protein-ligand H-bonds despite having two lone pairs and one proton, all of which can form H-bonds, can be rationalized by the resulting reduction in ligand selectivity. A ligand group with either good donor or acceptor geometry could both interact with that hydroxyl group, bringing the risk of misrecognition. This could have been the basis for negative selection during evolution.

**Table 1.** Intermolecular NH versus OH hydrogen bond donor frequencies for oxygen and nitrogen acceptors.

| H-bond donor molecule | H-bond type | Frequency | H-bond acceptor molecule |
| --- | --- | --- | --- |
| Protein | N-H ... O | 524 | Ligand |
| Protein | N-H ... N | 53 | Ligand |
| Protein | O-H ... O | 127 | Ligand |
| Protein | O-H ... N | 6 | Ligand |
| Ligand | N-H ... O | 219 | Protein |
| Ligand | N-H ... N | 1 | Protein |
| Ligand | O-H ... O | 124 | Protein |
| Ligand | O-H ... N | 1 | Protein |

### 2. Can the observed trends in interfacial polarity, with H-bonds tending to be formed by donors on the protein side of the interface interacting with acceptors on the ligand side, be explained by the prevalence of binding-site protons versus lone pairs?

To answer this question, the binding site was defined as all protein residues with at least one heavy atom within 9 Å of a ligand heavy atom. This set of potentially interacting atoms is typically used for interfacial analysis or scoring. All the previously mentioned criteria were then applied to identify intermolecular H-bonds, namely, meeting the 2.4-3.5 Å range for donor-acceptor distance and satisfying both the donor-H-acceptor and preacceptor-acceptor-H angular criteria. An example binding site and intermolecular H-bond network for one of the complexes appears in Fig. 4. For each binding site or ligand atom with H-bonding potential, the number of protons available to donate and the number of lone pairs available to accept H-bonds were tabulated and summed over the 136 complexes. The results (Fig. 5) show that acceptor lone pairs are significantly more prevalent than donor protons in the ligand binding sites of proteins (~16,000 lone pairs: ~10,000 protons available to donate), with a similar excess of lone pairs found in the ligands (~15,000 lone pairs: ~9,000 donor protons). Thus, if formation of intermolecular H-bonds were primarily driven by the prevalence of protons and lone pairs, the protein would be expected to accept H-bonds 1.6 times more often than it donates them. Given that the observed trend is in the opposite direction (a 2:1 tendency to donate H-bonds to the ligand; Fig. 3), there appears to be an underlying strong chemical or evolutionary preference for proteins to act as donors when binding cognate ligands.

**Fig. 4.**
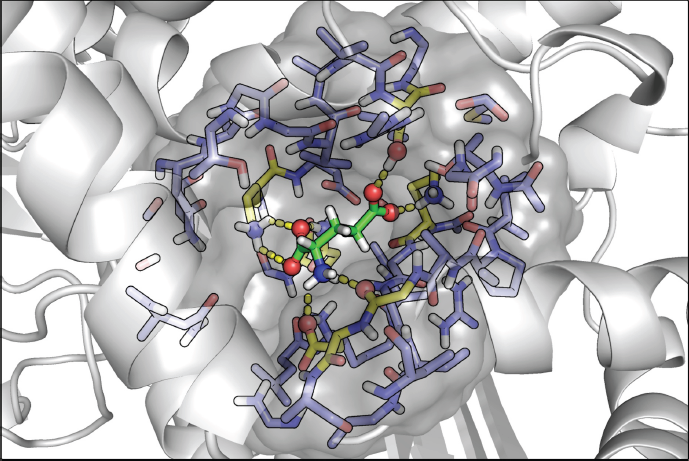
Binding site definition for glutamate hydrogenase interacting with glutamic acid ligand (PDB entry 1bgv). The gray solvent-accessible molecular surface envelops the ligand binding pocket defined as all protein atoms within 9 Å of the ligand’s heavy atoms (green tubes). The binding site residues H-bonding to the ligand are shown with carbon atoms in yellow, and all other binding site residues’ carbon atoms are colored in purple. Protein-ligand H-bonds as defined by Hbind are shown as yellow dashed lines.

**Fig. 5.**
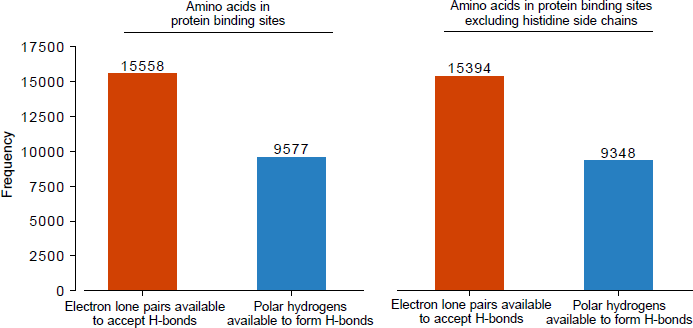
Statistics across 136 non-homologous complexes of the number of electron lone pairs in the protein’s binding site available to act as H-bond acceptors compared with the number or protons available to be donated. The observed frequencies indicate that ligand binding sites have a significant excess of lone pairs relative to protons that can participate in H-bonds. The analysis was performed with histidine side chains (two bars at left) and without (two bars at right), because this residue’s protonation state is more difficult to define. However, the histidine residues in the 136 complexes are primarily involved in metal interactions (in which the nitrogen lone pairs form bonds with cationic metals). Consequently, the statistics are substantially similar with and without histidine.

### 3. Do certain residues predominate in the observed preference for proteins to donate H-bonds to ligands?

The statistics of donor and acceptor atoms participating in interfacial H-bonds (Fig. 3) were further analyzed by atom type (Fig. 6). Panel (a) shows that amines, especially the terminal NH groups in Arg, Asn, Gln, and Lys, are the dominant donors of H-bonds to ligands, relative to hydroxyl groups. This cannot be explained by their prevalence in the binding sites. When the number of H-bonds formed is divided by the number of binding site occurrences, the H-bonding of terminal amines, especially in lysine, only becomes more pronounced (Fig. 6b). This is interesting, because Lys pays a higher entropic cost in lost degrees of bond-rotational freedom when H-bonding to ligands (due to having 4 side chain single bonds), relative to Arg (3 side chain single bonds) and especially Ser or Thr (2 single bonds). Lys, Ser and Thr can each potentially form up to three H-bonds with ligands, relative to Arg, which can form up to five. This also does not explain the preference for Lys. It could be that the greater flexibility and length of Lys and its rotatable proton positions (relative to the rigid and planar guanidinium group in arginine) allow lysine to better optimize H-bonds with ligands. Overall, the most prevalent H-bond donors and acceptors to ligands are the charge-bearing side chain atoms in Arg, Asp, Glu, and Lys, followed by the polar amine groups in Asn and Gln. The Asn and Gln NH_2_ groups form about 3 times as many ligand H-bonds as their terminal keto oxygens, despite having the capacity to form the same number of H-bonds per group. The same trend was observed when the number of interfacial H-bonds (Fig. 6a) was divided by the number of occurrences of each amino acid in the protein (Supplementary Material 5, Fig. S1) instead of the number of binding site occurrences (Fig. 6b). Interestingly, the same preference for very polar donors (Arg, Lys, and main-chain nitrogens) to participate in H-bonding with very polar acceptors (carboxylate and keto groups) has been found in analyses of both water-mediated and direct hydrogen bonds in protein-protein interfaces [29–31].

**Fig. 6.**
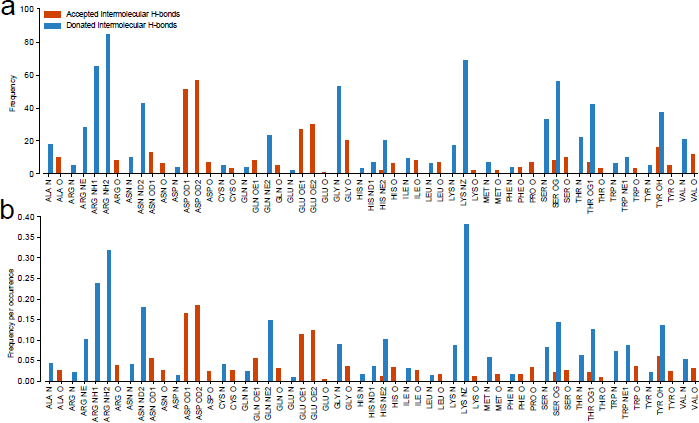
Intermolecular H-bonds formed by each amino acid atom type in ligand binding sites. (a) The frequency of H-bonds to ligand by atom type in 136 protein complexes. (b) The frequency per binding site occurrence of H-bonds to ligand. Pro N is omitted, because it lacks an amide proton to donate.

### 4. When protein and ligand atoms are categorized according to their chemistry, are H-bonding preferences between proteins and ligands fundamentally similar or different?

Protein atoms forming H-bonds with ligands were divided into main chain versus side chain categories (Fig. 7), and their H-bonds were tabulated according to atomic chemistry for keto oxygens (O), hydroxyl groups (OH), carboxylate oxygens (COO-), and amine nitrogens (NH and NH_2_). Amine donors were found to dominate in the total number of H-bonds formed with ligands, with almost equal representation from main and side chain amines (Fig. 7a). However, when normalized by the number of binding site occurrences, side chain amines were found to form 16 times as many ligand H-bonds as main chain amines (Fig. 7b). Hydroxyl groups donate a meaningful, though lesser, number of H-bonds to ligands (about one-fourth as many as amine groups donate) and rarely act as acceptors.

**Fig. 7.**
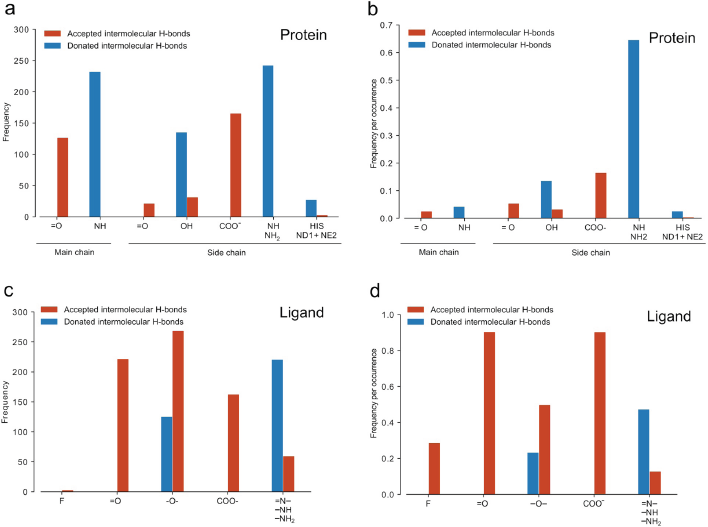
Comparison of the chemistry and prevalence of atoms forming intermolecular H-bonds, by protein versus ligand side of the interface. The bar plot in (a) shows the frequency of protein atoms participating in H-bonds to ligands in the 136 complexes, while (b) shows the same data normalized by the number of binding site occurrences, yielding the average number of H-bonds to ligand per atom type. The bar plots (c) and (d) show the same data from the ligands' perspective. In panels (c) and (d), F indicates fluorine. The label =O includes O.2 (sp2-hybridized) oxygen atoms; −O- includes O.3 (sp3-hybridized) hydroxyl and ester oxygens; COO- includes O.co2 oxygen atoms in carboxylate, sulfate, and phosphate groups; and =N, −NH, and – NH_2_ includes N.2, N.3, N.am, N.ar. For COO- groups, each terminal oxygen was tabulated separately.

Surprisingly, the trends for H-bond donors and acceptor chemistry in ligands are quite different (Fig. 7c and 7d). Keto (=O), ester + hydroxyl (–O–), and carboxylate oxygens dominate the total number of H-bonds formed with proteins (Fig. 7c). When the number of H-bonds formed is normalized by the number of occurrences of each atom type in the ligands, it becomes apparent that on average 90% of all keto and carboxylate oxygens in biological ligands form direct H-bonds to their proteins. Ligand fluorine and nitrogen atoms also participate as H-bond acceptors. Consistent with the observed strong trend for ligands to accept rather than donate H-bonds to proteins, ligand hydroxyl and amine donors only form one-third as many protein H-bonds per occurrence when compared with oxygen acceptors.

These results are also consistent with results from an earlier analysis of water molecules forming H-bonded bridges between proteins and ligands (A. Cayemberg and L. A. Kuhn, unpublished results) in a set of 20 non-homologous complexes [32]. There, without defining donor or acceptor roles, we discovered that water molecules H-bonding directly to both the protein and ligand interacted with oxygen atoms on the ligand 74% of the time and nitrogen atoms only 25% of the time (with Cl atoms representing the final 1%). The same was true for water molecules forming di-water bridges between protein and ligand, with a 76% preference for interacting with oxygen on the ligand. Hong & Kim (2016) [29] noted that interfacial water molecules, on average, make more interactions with protein H-bond acceptors than donors, similar to our finding that protein-bound water molecules interact with ligand oxygen atoms (primarily H-bond acceptors) more often than nitrogen atoms (primarily donors).

### 5. Do different classes of ligand differ in their tendency to accept versus donate H-bonds?

For this analysis, the 136 complexes were considered from the ligand perspective, with the 25 peptidyl, 50 nucleotide-like, and 61 other small organic ligands analyzed as individual sets (Supplementary Material 1, Table S1). The 2:1 ratio for ligands to accept rather than donate H-bonds to cognate proteins was seen for both nucleotidyl and other organic ligands (Fig. 8). Peptidyl ligands, on the other hand, showed no strong preference for donating versus accepting H-bonds. This is expected, because of the fundamental chemical and evolutionary parity between the peptides and proteins in these complexes: both cannot act primarily as donors and still make sufficient intermolecular H-bonds. The more polar, often charged, nucleotidyl ligands formed 50% more H-bonds with proteins than the other organic molecules. This is in line with the observation (Fig. 6) that charged protein side chains play a more important role than neutral side chains in H-bonding to ligands. The strength of an H-bond also increases with the magnitude of the complementary charge on the participating atoms [33]. However, the greater number of H-bonds for nucleotidyl ligands could also reflect their greater number of heavy atoms, 31.7 +/- 11.2 on average, relative to other organic molecules, 17.8 +/- 10.8. The average number of heavy atoms for peptidyl ligands was 27.8 +/- 21.1.

**Fig. 8.**
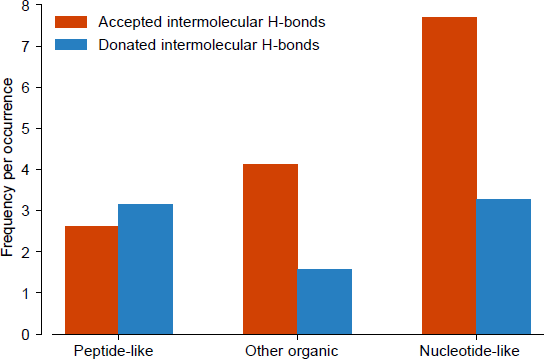
The average number of H-bonds donated (blue) or accepted (red) for each ligand type: peptidyl, nucleotide-like, and other organic.

These results indicate that strong H-bonds involving charged groups are common in cognate protein-ligand complexes. The prevalence of strong H-bonds involving very polar groups is not necessarily expected, given that ligands need to be released from their proteins as part of the enzymatic, signaling, or transport cycles. Strong H-bonds also contribute to formation of the catalytic transition state between enzymes and their ligands [33].

Given the visual observation of dense protein networks of nitrogen H-bond donors interacting with the nucleotides, nucleotidyl ligands were the only ligand class for which a clear, local pattern of H-bonding appeared, involving at least 3 nitrogen H-bond donor groups separated by no more than 10 residues (Fig. 9). Three to five H-bond donors occurred within six residues of the amino acid sequence (positions 1-6 in Fig. 9) in 18 of the 24 cases (rows 2-19). Thirteen of the 18 patterns involved nucleotidyl ligands, with the sequence pattern **Gly-Lys-(Ser,Thr)-(Thr,Ser,Tyr**,Cys,Ala) found in 7 cases. (Boldface indicates the dominant residue type(s) and regular font indicates other allowed residues.) This structural motif turns out to be the P-loop nest for phosphate binding, a strong and particularly geometrically ordered example of the tendency for proteins to donate H-bonds to ligands [34] (Fig. 10). The program for creating PyMOL H-bond interaction views from Hbind tables, as shown in this figure, is freely available to researchers at https://github.com/psa-lab/HbindViz.

**Fig. 9.**
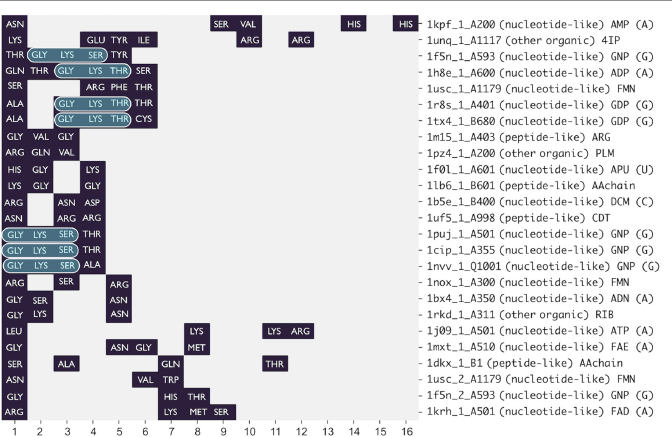
Patterns of H-bonds to ligands that are localized in the protein sequence and involve three or more nitrogen donors. The x-axis indexes from the first to the sixteenth position in all amino acid sequences with no more than 10 residues between adjacent H-bond donors to ligand. The label in the rightmost column provides the PDB code and index of the H-bond pattern (1, 2, etc.) in a given protein, the chain ID and residue number of the ligand in the PDB structure file, the ligand category (nucleotide-like, peptidyl, or other organic), and the 3-letter ligand name in the PDB. Where appropriate, the base (adenine, A; guanine, G; or C, deoxycytidine) present in the nucleotide-like ligands is provided at the end of the label. Highlighted in blue are the Gly-Lys-Ser/Thr motifs found hydrogen-bonding to phosphate groups in seven of the nucleotidyl ligands.

**Fig. 10.**
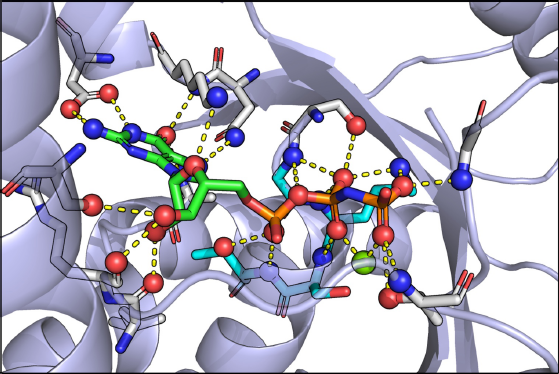
P-loop nest motif Gly-Lys-Ser-Thr for phosphate binding, with carbon atoms in cyan, donating a local network of protein H-bonds to the oxygen-rich triphosphate group. This example is from a high-resolution G protein structure in complex with GTP (PDB entry 1cip [45]). H-bonds forming the P-loop nest interaction are shown as yellow dashed lines, and polar atoms participating in these interfacial interactions appear as red spheres for oxygen atoms, blue spheres for nitrogen atoms, and a green sphere for the bound Mg^2+^. For clarity, hydrogen atoms are omitted.

### 6. Can orientational selectivity of the biological ligand explain the preference for proteins to donate H-bonds to ligands?

Here we evaluate whether geometrical aspects of H-bond interactions, in particular the angular dependence of H-bonds, can provide a ligand-selectivity advantage in proteins that donate H-bonds to ligands more often than accepting them. Underlying protein-ligand binding is a 3D code defined by structure and chemistry that determines which ligands can bind to a protein, as well as avoiding binding to ligands that inappropriately alter activity.

An interesting example of how strong selectivity for a molecular partner can confer a functional advantage is the observation that narrow-spectrum (more selective) antibiotic ligands avoid drug resistance much more effectively than broad-spectrum antibiotics [35]. Narrow-spectrum antibiotics form interactions that are highly tuned for their protein target, which means that the protein must accumulate more mutations to abrogate binding by a narrow-spectrum antibiotic, relative to broad-spectrum antibiotics. The same effect, in the absence of any mutations, allows proteins that form many ligand-selective interactions to prevent misrecognition and binding to the wrong partners.

The simplest case supporting a hypothesized preference for proteins to use more chemically or geometrically selective interactions in ligand binding is the observed 3:1 preference for proteins to use amines relative to hydroxyl groups in ligand H-bonds (797 amine-involving H-bonds versus 258 hydroxyl-involving H-bonds; Table 1). This is despite the potential of the hydroxyl group to accept two H-bonds and donate one, which would allow about 1.5 times as many H-bonds to the ligand relative to the most common protein amine groups (NH and NH_2_). In general, protein lone pairs available to accept H-bonds are 1.6 times as prevalent as protons available to donate (Fig. 5). However, the hydroxyl group is less selective in its interactions, allowing both donor and acceptor groups on ligand partners, which may result in insufficient selectivity for the correct ligand relative to the thousands of alternative molecules in the cell.

Ligand selectivity can also be conferred by the difference in geometrical constraints on donor versus acceptor interactions. To examine how selectivity relates to the 3D geometry of interaction, the favored angular and donor-acceptor distance ranges are shown for H-bond acceptor and donor atoms (Fig. 1). A favorable donor–H···acceptor angle Θ range of 120–180˚ in well-resolved crystal structures, in combination with a favorable donor–acceptor separation of 2.4-3.5 Å [6, 22], results in a significantly smaller volume (30.3 Å^3^) in which a ligand acceptor atom can favorably interact with a protein donor atom, in comparison with the volume in which a ligand proton can favorably interact with lone pairs on a protein acceptor (60.6 Å^3^). This is partly due to the more permissive pre-acceptor–acceptor–H angle (ϕ) of 90-180˚ (relative to the Θ constraint on donor–H…acceptor angle), and also due to the presence of two lone pairs on the majority of H-bond acceptor atoms in proteins (oxygens). The two lone pairs create a large, continuous volume in which a proton can H-bond with the acceptor atom. The observed distribution of donor atoms relative to oxygen acceptor atoms in well-resolved protein X-ray structures [6] indicates there are few constraints on out-of-plane interactions with acceptor lone pairs, resulting in an almost isotropic, hemispheric shell of donor proton positions relative to the acceptor.

An evolutionary emphasis on matching large volumes of favorable interaction around acceptors on the protein might well result in too little selectivity for the cognate ligand. While proteins, with their current amino acid content, cannot avoid the presence of oxygen atoms on the surface, nor do proteins entirely avoid ligand interactions with acceptors, we hypothesize that cognate protein-ligand interactions may have evolved to favor the use of donor groups on the protein to create small volumes that the arrangement of acceptor atoms on cognate ligands must uniquely match. This is supported by the enhancement of oxygen atoms on small molecule ligands (Fig. 7c). It is also supported by an observation of Taylor et al (1984) [36]: though the majority of intramolecular N–H**…**O H-bond angles are in the 100–140˚ range, intermolecular N–H**…**O angles are typically much more linear (170–180˚), corresponding to stronger H-bonds as well as a narrow tolerance to be met in recognizing the cognate ligand. Donation of H-bonds to the ligand is, of course, one component of recognition. Shape complementarity, hydrophobic surface matching, interfacial ion binding, and additional H-bonds including water-mediated interactions [32, 36–38] complete the selection of and enhanced affinity for the native ligand.

Another selective advantage that could drive the evolution of strong donor patterns (rather than mixed donor-acceptor patterns) for ligand binding, is to disfavor aberrant protein-protein interaction. Binding site donor geometries that evolved to match a small molecule ligand could not easily be satisfied by other proteins, which on average also favor binding site donor patterns that would tend to repel interaction with other sets of donors. Finally, the finding that asymmetry in packing of the peptide amide dipole results in larger positive than negative regions in proteins [39] would tend to enhance the preference for proteins to interact with more electronegative, lone-pair bearing atoms.

Another factor that comes into play in the observed preference for biological ligands to accept hydrogen bonds from proteins is their enhancement in oxygen atoms relative to nitrogen. Within cells, oxygen (particularly derived from water) is available at higher concentration than nitrogen, and therefore oxygen-rich ligands may be more readily synthesized. Oxygen is also more abundant in solid earth, in fact 10,000 times more abundant than nitrogen, due to the inability of nitrogen to contribute to stable lattices [40], though nitrogen is more abundant in the atmosphere. Plants are the source of nutrients for many other organisms, and they can more readily incorporate environmental oxygen from CO_2_ into metabolites, relative to N_2_, which requires fixation. To consider their abundance within biological ligands, we compared the number of oxygen versus nitrogen atoms in the 136 ligands in our study and also in 96 metabolites characterized in *Mycoplasma genitalium*, one of the simplest organisms [41]. Oxygen atoms were found to be 2.6 times as prevalent as nitrogen atoms in the *M. genitalium* metabolites and 2.1 times as prevalent as nitrogen in the 136 ligands in the current study. Of course, the two-fold preference for hydrogen bond accepting by ligand atoms (mainly oxygen atoms; Fig. 7C) does not only reflect the prevalence of oxygen atoms; the preference for accepting H-bonds when normalized by atom type is even stronger (Fig. 7D). Still, it is quite interesting, and not widely recognized, that oxygen atoms are much more common than nitrogen in biological ligands and that their acceptor role is so dominant. Both features can be useful in designing new ligands that successfully compete for binding.

### 7. Do protein-bound metal ions contribute significantly to ligand binding, and how does their bond chemistry relate to observed trends in H-bonding?

When protein-bound metal ions were found in the ligand interface, they were included in the analysis. Table S2 (Supplementary Material 6) provides detailed statistics, while Fig. 11 summarizes ligand interactions per occurrence for the 8 metal types observed in the 136 complexes. Mg^2+^ was by far the most common, with 24 occurrences, followed by Mn^2+^ with 14 occurrences. All other metal types were present 7 or fewer times. Ni^2+^, Mg^2+^, Cd^2+^, Mn^2+^, Co^2+^, and Na^+^ each accounted for 1-2 direct ligand bonds per occurrence (using bond-length criteria listed in Methods), while Fe (exhibiting various oxidation states in the different complexes) and Zn^2+^ averaged half an interaction per occurrence. Metal interactions with lone pairs on electronegative atoms within bonding distance, as measured here, are almost covalent in strength. This makes them significant contributors to the enthalpy change upon complex formation. Because these metals are positively charged, the trend in polarity of the interface is like the dominant H-bond classes observed above, with a positively charged group on the protein side forming a bond with a lone pair of electrons on the ligand.

**Fig. 11.**
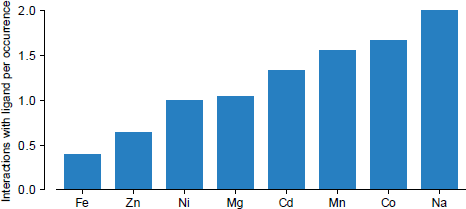
The average number of bonds to ligand formed per occurrence by protein-bound metal ions in the 136 complexes.

### 8. How do these results relate to Lipinski’s rule of 5 for drug-likeness in small molecules?

In the late 1990s, Lipinski and colleagues at Pfizer undertook a study of 2,245 small molecules from the World Drug Index considered to have superior physicochemical properties, based on meeting solubility and cell permeability criteria required for entry into Phase II clinical trials [8]. The drug-like criteria for small molecules derived from their analysis of the 2,245 compounds are known as the Rule of 5. Poor absorption or permeability tends to occur for a compound matching any of the Rule of 5 features: more than 5 H-bond donors, more than 10 H-bond acceptors, a molecular weight greater than 500 Da, or a calculated logP value of greater than 5, defined as the logarithm of the partition coefficient between n-octanol and water. These criteria remain widely used for selecting sets of molecules for virtual or high-throughput experimental screening, and as ideal physicochemical ranges to match when redesigning lead compounds to bind with higher affinity or better bioavailability. The Rule of 5 criteria were not derived to predict molecules as effective protein ligands. However, most drugs do target proteins, and thus the Rule of 5 criteria may select for the ability to bind proteins as well as enter the cell. In fact, the maximum H-bond acceptor to donor ratio in the Rule of 5 (10:5) matches the trend found here: twice as many H-bonds being accepted by ligands (5 on average) as donated (2.5 on average; Fig. 3). The two-fold preference for ligand acceptors relative to donors in H-bonding may therefore be a molecular mechanism underlying the drug-like criteria in the Rule of 5. Additionally, the ability of H-bond acceptor and donor numbers to predict drug-likeness suggests that the trends identified in this paper can also be useful predictors.

### 9. Can the observed H-bonding trends be used to predict protein-ligand interactions?

We addressed the question of whether the observed prevalence of H-bond acceptors and donors in the 136 complexes, tabulated by PDB atom type for protein binding site atoms (e.g., Arg O, N, NE, NH1, and NH2) and by MOL2 atom type for ligand atoms (e.g., O.2, O.3, N.2, N.3, etc.), can be used to predict the cognate protein-ligand orientation from a series of dockings of the small molecule. To test this, we used 10 ligand dockings on average in each of 30 protein-small molecule complexes that were recently used in a comparison of docking scoring functions. This set does not overlap with the 136 complexes [42] (Supplementary Material 4, Table S3). The crystallographic binding pose was not included, because the correct pose is unknown in a predictive study and therefore never exactly sampled. Secondly, many scoring methods can readily detect the crystallographic pose as the global optimum due to their parameterization, suggesting excellent accuracy when the crystal pose is included; a much more realistic assessment of their real-world performance is the identification of near-native poses. The best-sampled ligand docking poses here ranged from 0.1-1.4 Å RMSD relative to the crystallographic position across the 30 complexes, as shown by the green cumulative distribution curve in Fig. 12. The goal of this analysis of docked positions was not to develop a new scoring function, but to assess whether the H-bond interaction statistics accumulated across 136 structures capture the essential molecular recognition features that occur within individual structures sufficiently well to discriminate native or near-native interactions.

**Fig. 12.**
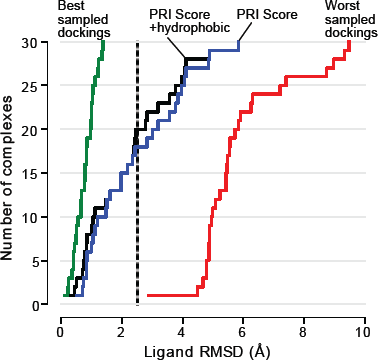
Enrichment plot showing the degree of native-likeness (RMSD relative to crystallographic position) of docking showing the highest Protein Recognition Index for all 30 complexes. The ligand orientation for each complex was predicted according to the highest H-bond PRI value (blue trace) or the highest PRI+ hydrophobic contact score (black trace) among all the ligand orientations. All 30 complexes’ best-scoring ligand orientations were then compiled, and their RMSD values relative to the crystallographic position were sorted from best (closest to 0 Å) to worst RMSD (4-5 Å). These RMSD values were then plotted as a cumulative distribution function of the number of ligand orientations selected to within X Å RMSD of the crystallographic position. For instance, all ligand orientations selected by either PRI or PRI + hydrophobic scoring that appear to the left of the dashed black vertical line at ligand RMSD = 2.5 Å were within 2.5 Å RMSD of the crystallographic position. This was true for 18 of the PRI scored complexes and 20 of the PRI + hydrophobic scored complexes. The result that would be obtained by the best-possible scoring of ligand orientations (selecting the best-sampled docking of the ligand for each complex) is shown by the green trace. The result from selecting the worst docking (highest RMSD position) of each ligand across the 30 complexes is shown by the red trace.

For the protein H-bond component of the scoring function, the frequency scores of all protein atoms observed to make an H-bond with the ligand were summed, based on the raw data compiled across the 136 complexes, with sample data shown below. In the first entry, {Acceptor: 0, Donor: 18} indicates that in the 136 complexes, alanine main chain nitrogen atoms accepted H-bonds from the ligand 0 times and donated H-bonds to ligands 18 times.

ALA:
N: {Acceptor: 0, Donor: 18}
O: {Acceptor: 10, Donor: 0}

ARG:
N: {Acceptor: 0, Donor: 5}
NE: {Acceptor: 0, Donor: 28}
NH1: {Acceptor: 0, Donor: 65}

*etc.*

So, for instance, if you were to score a ligand orientation accepting an H-bond from the main chain N in Ala and H-bonds from both Arg NE and Arg NH1, the protein H-bond score for that binding mode would be:

18 + 28 + 65 = 111

The higher the score, the more the docking reflects the known preferences in the 136 complexes for H-bonds donated or accepted by the protein. This Protein Recognition Index, or PRI-prot, differs from the typical scoring of H-bonds in protein-ligand docking, because here the contribution of each H-bond is weighted according to the prevalence of intermolecular H-bonds involving this protein atom type in crystal complexes. Scoring is performed the same way for the ligand side of the interaction, leading to a PRI-lig value. Standardization is then performed on the PRI-lig values across the dockings for a given complex, rescaling such that the score distribution has a mean value of 0 and a variance of 1. This converts the PRI-lig to a Z-score measured in standard deviations above or below the mean (more favorable or less favorable, statistically). The same standardization is performed for protein PRI-prot values across the dockings, putting the ligand and protein PRI values on the same scale. PRI-prot and PRI-lig values can then be summed (reflecting the simplest possible weighting, giving equal importance to the protein and ligand side of the interface), to yield what we call the PRI. High PRI values reflect that the H-bond groups linked between protein and ligand in the current ligand orientation match the H-bond preferences found in the 136 unrelated complexes. For a series of ligand dockings in a given protein, the docking with the highest PRI is predicted as the most native-like complex. This process was performed for all 30 complexes, and the results are summarized in Fig. 12. To consider the extent to which hydrophobic contacts add information for defining the cognate ligand orientation, we created a variant of PRI that includes an equal-weighted, standardized hydrophobic contact term (PRI+hydrophobic). The hydrophobic term counts the number of carbon-carbon and carbon-sulfur contacts (atom centers within 4 Å) between the protein and ligand, as reported in the Hbind software output (Fig. 2). Software we used to compute the Protein Recognition Index (and its PRI-prot and PRI-lig) components is available at https://github.com/psa-lab/protein-recognition-index. We envision this software will be a broadly useful tool for assessing the native-likeness of designed or predicted protein-ligand interfaces, as well as for guiding protein mutagenesis to identify ligand binding residues and predict ligand binding sites (using the PRI-prot component alone) or assessing ligand physicochemical suitability for a protein target (using the PRI-lig component alone).

The results of the ligand orientation prediction enrichment plot (Fig. 12) clearly show that the statistical information encoded in the Hbind hydrogen-bonding preferences of different atom types is able to identify near-native ligand orientations, selecting an orientation within 2.5 Å RMSD of the crystallographic position in two-thirds of the complexes. Adding a hydrophobic contact term leads to a slight improvement in prediction, while the H-bonding preferences account for most of the predictive power. Measuring the Pearson linear correlation coefficient (r) between the PRI values, PRI+hydrophobic values, and two commonly used docking scoring functions, AutoDock Vina [43] (version 1.1.2; http://vina.scripps.edu) and DSX [44] (also known as DrugScore X; version 0.88; http://pc1664.pharmazie.unimarburg.de/drugscore/) across ~300 dockings for the 30 complexes, shows that the PRI value is almost uncorrelated with the scores from AutoDock Vina (r = −0.26) and DSX (r = −0.19), despite these scoring functions also including H-bond interaction terms. This indicates that PRI provides new information that has high predictive value on its own, while also easily being combined with existing protein-ligand scoring metrics. Weighting H-bonds according to their statistical prevalence by atom type measures a chemical aspect of protein-ligand recognition that is both predictive of native interactions and not reflected in the other measures.

## Conclusions

To address the question that motivated this work — whether proteins tend to donate rather than accept H-bonds when binding biological small molecules — a utility called Hbind was developed to label the donor/acceptor capacity of each atom, and characterize each H-bond in terms of its atomic chemistry and geometry. Making this software available allows such data to be generated readily and analyzed for a range of other interesting questions with the vast crystal structure data now available. Handling both protein and ligand chemistry at the atomic rather than coarser functional group or side chain levels allowed an in-depth analysis of the trends and potential underlying mechanisms in ligand recognition by proteins. Our conclusions are:

- Across 136 non-homologous protein complexes including a mix of nucleotide-like, peptidyl, and other organic ligands, the proteins were found to donate twice as many H-bonds as they accepted from ligands.
- Lone pairs available to accept H-bonds are actually 1.6 times as prevalent as protons available to donate, both on the protein and ligand side of the interface. Thus, the relative availability of donor and acceptor groups does not explain the trend for proteins to preferentially donate H-bonds to their ligands.
- A corresponding, strong preference for ligands to accept H-bonds from proteins suggests that focusing on the prevalence and positioning of H-bond acceptors in both designed ligands and molecules assessed in screening (that is, a more detailed, structural measure of “drug-likeness”) is likely to result in ligands that better match the protein-encoded determinants for binding. The Protein Recognition Index (PRI) software was designed for this purpose.
- Furthermore, on average 90% of all keto and carboxylate oxygens in the biological ligands were found to form direct hydrogen bonds to their protein partners. This suggests that satisfying these oxygen atoms with direct protein H-bonds is an important and unrecognized feature of native ligand recognition.
- Nitrogen atoms served as donors for 76% of the intermolecular H-bonds and hydroxyl groups in 24%, considering both protein and ligand donors together. This suggests that amine nitrogen atoms are much more effective donors in biological complexes than hydroxyl groups, providing another straightforward way to enhance molecular design.
- The side chains in proteins most likely to donate H-bonds to ligands are Arg and Lys, with Asn and Gln being about half as important. Asp and Glu are the side chains most likely to accept H-bonds from ligands. Highly polar H-bonds are apparently favored in the underlying code of molecular recognition. These results suggest focusing on these side chains when predicting binding sites or carrying out experiments to identify key H-bonding groups within a site.
- Metals bound in protein ligand-binding sites are not a dominant feature. Most metal ions in binding sites account for 1-1.7 bonds to the ligand, on average, with Fe and Zn accounting for fewer ligand interactions (0.5, on average) in the 136 complexes. While these bonds occur less frequently, their almost-covalent strength makes them important contributors to affinity.
- These trends, analyzed from all angles, indicate a surprising degree of interfacial polarization for non-peptidyl organic molecule complexes with proteins, favoring donors on the protein side and acceptors on the ligand side. The pairing of amine donors on the protein with oxygen acceptors on the ligand is a dominant motif in protein-small molecule recognition, like the amine-keto pairing between bases in nucleic acids.
- By developing software to calculate a Protein Recognition Index (PRI), measuring the similarity between H-bonding features in a given complex (predicted or designed) and the characteristic H-bond trends from crystallographic complexes (Fig. 7), we show that the cognate orientation between protein and ligand can be predicted from this information alone. The PRI for a set of protein or ligand atoms can also be calculated, to discern the extent to which their H-bonding groups match the favored distribution of donor and acceptor atom types in known complexes.
- The 2:1 acceptor to donor ratio observed here for ligand atoms forming H-bonds to proteins appears to be an underlying structural explanation for the 2:1 ratio of ligand H-bond acceptor atoms to donor atoms in Lipinski’s Rule of 5. We anticipate the Protein Recognition Index may prove similarly useful in guiding protein and ligand design to design more selective and tighter-binding complexes.
- The trend for proteins to donate H-bonds to their cognate ligands, especially via amine donor groups, may have evolved as a ligand selectivity determinant. Amine donors have relatively narrow angular constraints and volumes in which an acceptor group can form an energetically favorable H-bond. Two acceptor lone pairs are present on the oxygen atoms in proteins, and a consequence is that the lone pairs present a broad surface and volume for favorable interaction with donor atoms (twice that of an NH donor interacting with an acceptor group). Molecular evolution is expected to favor a narrow selection of ligand partners due to the potential for misrecognition if many ligands could easily match H-bonding groups in a protein pocket. The relative orientation and spacing of these groups is also an extremely important aspect of the code for matching H-bonds between protein and ligand.

## Acknowledgments

This research was supported by funding from the Great Lakes Fishery Commission (Project ID: 2015_KUH_54031). We gratefully acknowledge OpenEye Scientific Software (Santa Fe, NM) for providing academic licenses for the use of their QUACPAC (molcharge) and OEChem software. We also thank the following lab graduates for their contributions to this research: Dr. Maria Zavodszky (now at GE Global Research Center), who observed that hydroxyl-rich ligands tended to result in false positives in screening, Dr. Amy Cayemberg McQuade (now at Carroll University) for carrying out the statistical analysis of protein-water-ligand hydrogen-bond bridges, and Dr. Jeffrey VanVoorst (now at Veritas Technologies, LLC) for developing the non-homologous dataset of 136 protein-small molecule complexes analyzed here. We thank Michael Feig (Michigan State University) for discussions on the biological basis for the prevalence of oxygen versus nitrogen in natural ligands and also appreciate the data he provided on the atomic composition of metabolites in *Mycoplasma genitalium*.

## Supplementary Materials

- Table of 136 complexes analyzed (Supplementary Material 1, Table S1; supp_table_S1.xlsx)
- YASARA protonation script (Supplementary Material 2, supp_material_2_yasara_opthyd_macro.txt)
- MOL2 structure files for all 136 ligands as prepared (Supplementary Material 3, supp_material_3_lig_in_orig_PDB_refframe.mol2)
- Hbind interaction files for all 136 complexes (Supplementary Material 4, supp_material_4_hbind_interaction_tables.txt)
- Bar plot showing the number of H-bonds by amino acid atom type normalized by the number of the atom type occurrences in proteins (Supplementary Material 5, Fig. S1; supp_material_5_fig-S1.pdf)
- Table of metal interaction statistics (Supplementary Material 6, Table S2, supplementary_table_S2.xlsx)
- Table of 30 protein complexes for analyzing PRI (Supplementary Material 7, Table S3, supp_material_7_table_S3.xlsx)

## References

1. Zavodszky MI, Sanschagrin PC, Korde RS, Kuhn LA (2002) Distilling the essential features of a protein surface for improving protein-ligand docking, scoring, and virtual screening. J Comput Aided Mol Des 16:883–902.

2. Sukuru SCK, Crepin T, Milev Y, Marsh LC, Hill JB, Anderson RJ, Morris JC, Rohatgi A, O’Mahony G, Grøtli M, others (2006) Discovering new classes of Brugia malayi asparaginyl-tRNA synthetase inhibitors and relating specificity to conformational change. J Comput Aided Mol Des 20:159–178.

3. Zavodszky MI, Rohatgi A, Van Voorst JR, Yan H, Kuhn LA (2009) Scoring ligand similarity in structure-based virtual screening. J Mol Recognit 22:280–292.

4. Van Voorst JR, Tong Y, Kuhn LA (2012) ArtSurf: a method for deformable partial matching of protein small-molecule binding sites. In: Proc. ACM Conf. Bioinformatics, Comput. Biol. Biomed. pp 36–43

5. Nittinger E, Inhester T, Bietz S, Meyder A, Schomburg KT, Lange G, Klein R, Rarey M (2017) Large-scale analysis of hydrogen bond interaction patterns in protein-ligand interfaces. J Med Chem 60:4245–4257.

6. McDonald I, Thornton JM (1994) Atlas of side-chain and main-chain hydrogen bonding. Biochemistry and Molecular Biology Department, University College London, http://www.biochem.ucl.ac.uk/bsm/atlas

7. Panigrahi SK, Desiraju GR (2007) Strong and weak hydrogen bonds in the protein–ligand interface. Proteins Struct Funct Bioinforma 67:128–141.

8. Lipinski CA, Lombardo F, Dominy BW, Feeney PJ (1997) Experimental and computational approaches to estimate solubility and permeability in drug discovery and development settings. Adv Drug Deliv Rev 23:3–25.

9. Dawson NL, Lewis TE, Das S, Lees JG, Lee D, Ashford P, Orengo CA, Sillitoe I (2016) CATH: an expanded resource to predict protein function through structure and sequence. Nucleic Acids Res 45:289–295.

10. Ahmed A, Smith RD, Clark JJ, Dunbar Jr JB, Carlson HA (2014) Recent improvements to Binding MOAD: a resource for protein-ligand binding affinities and structures. Nucleic Acids Res 43:465–469.

11. Berman HM, Westbrook J, Feng Z, Gilliland G, Bhat TN, Weissig H, Shindyalov IN, Bourne PE (2006) The Protein Data Bank. In: Int. Tables Crystallogr. Vol. F Crystallogr. Biol. Macromol. Springer, pp 675–684

12. Warren GL, Do TD, Kelley BP, Nicholls A, Warren SD (2012) Essential considerations for using protein–ligand structures in drug discovery. Drug Discov Today 17:1270–1281.

13. Krieger E, Joo K, Lee J, Lee J, Raman S, Thompson J, Tyka M, Baker D, Karplus K (2009) Improving physical realism, stereochemistry, and side-chain accuracy in homology modeling: Four approaches that performed well in CASP8. Proteins Struct Funct Bioinforma 77:114–122.

14. Krieger E, Dunbrack RL, Hooft RWW, Krieger B (2012) Assignment of protonation states in proteins and ligands: combining pK a prediction with hydrogen bonding network optimization. Methods Mol Biol Comput Drug Discov Des 819:405–421.

15. Colominas C, Luque FJ, Orozco M (1996) Tautomerism and protonation of guanine and cytosine. Implications in the formation of hydrogen-bonded complexes. J Am Chem Soc 118:6811–6821.

16. Krieger E, Darden T, Nabuurs SB, Finkelstein A, Vriend G (2004) Making optimal use of empirical energy functions: force-field parameterization in crystal space. Proteins Struct Funct Bioinforma 57:678–683.

17. Jakalian A, Jack DB, Bayly CI (2002) Fast, efficient generation of high-quality atomic charges. AM1-BCC model: II. Parameterization and validation. J Comput Chem 23:1623–1641.

18. DeLano WL (2002) Pymol: An open-source molecular graphics tool. CCP4 Newsl Protein Crystallogr 40:82–92.

19. Halgren TA (1996) Merck molecular force field. I. Basis, form, scope, parameterization, and performance of MMFF94. J Comput Chem 17:490–519.

20. Tripos (2007) Tripos Mol2 File Format. St Louis, MO http://www.tripos.com/data/support/mol2.pdf.

21. Pauling L (1960) The nature of the chemical bond and the structure of molecules and crystals: an introduction to modern structural chemistry. Cornell University Press, Ithaca, New York, p 452

22. Ippolito JA, Alexander RS, Christianson DW (1990) Hydrogen bond stereochemistry in protein structure and function. J Mol Biol 215:457–471.

23. Prakash B, Renault L, Praefcke GJK, Herrmann C, Wittinghofer A (2000) Triphosphate structure of guanylate-binding protein 1 and implications for nucleotide binding and GTPase mechanism. EMBO J 19:4555–4564.

24. Van Der Walt S, Colbert SC, Varoquaux G (2011) The NumPy array: a structure for efficient numerical computation. Comput Sci Eng 13:22–30.

25. Jones E, Oliphant T, Peterson P (2001) SciPy: Open source scientific tools for Python. http://www.scipy.org

26. McKinney W (2010) Data structures for statistical computing in Python. In: Millman J, van der Walt S (eds) Proc. 9th Python Sci. Conf. pp 51–56

27. Raschka S (2017) BioPandas: Working with molecular structures in pandas DataFrames. J Open Source Softw 2:1–3.

28. Hunter JD (2007) Matplotlib: A 2D graphics environment. Comput Sci Eng 9:90–95.

29. Hong S, Kim D (2016) Interaction between bound water molecules and local protein structures: A statistical analysis of the hydrogen bond structures around bound water molecules. Proteins Struct Funct Bioinforma 84:43–51.

30. Miyazawa S, Jernigan RL (1996) Residue–residue potentials with a favorable contact pair term and an unfavorable high packing density term, for simulation and threading. J Mol Biol 256:623–644.

31. Glaser F, Steinberg DM, Vakser IA, Ben-Tal N (2001) Residue frequencies and pairing preferences at protein–protein interfaces. Proteins Struct Funct Bioinforma 43:89–102.

32. Raymer ML, Sanschagrin PC, Punch WF, Venkataraman S, Goodman ED, Kuhn LA (1997) Predicting conserved water-mediated and polar ligand interactions in proteins using a K-nearest-neighbors genetic algorithm. J Mol Biol 265:445–464.

33. Shan S, Herschlag D (1996) The change in hydrogen bond strength accompanying charge rearrangement: Implications for enzymatic catalysis. Proc Natl Acad Sci 93:14474–14479.

34. Bianchi A, Giorgi C, Ruzza P, Toniolo C, Milner-White EJ (2012) A synthetic hexapeptide designed to resemble a proteinaceous p-loop nest is shown to bind inorganic phosphate. Proteins Struct Funct Bioinforma 80:1418–1424.

35. Palumbi SR (2001) Humans as the world’s greatest evolutionary force. Science (80-) 293:1786–1790.

36. Taylor R, Kennard O (1984) Hydrogen-bond geometry in organic crystals. Acc Chem Res 17:320–326.

37. Craig L, Sanschagrin PC, Rozek A, Lackie S, Kuhn LA, Scott JK (1998) The role of structure in antibody cross-reactivity between peptides and folded proteins. J Mol Biol 281:183–201.

38. Kuhn LA, Swanson CA, Pique ME, Tainer JA, Getzoff ED (1995) Atomic and residue hydrophilicity in the context of folded protein structures. Proteins Struct Funct Bioinforma 23:536–547.

39. Gunner MR, Saleh MA, Cross E, Wise M, others (2000) Backbone dipoles generate positive potentials in all proteins: origins and implications of the effect. Biophys J 78:1126–1144.

40. Rubin K. Ask an Earth Scientist. https://www.soest.hawaii.edu/GG/ASK/atmonitrogen.html. (Date accessed: 01/17/2018).

41. Feig M, Harada R, Mori T, Yu I, Takahashi K, Sugita Y (2015) Complete atomistic model of a bacterial cytoplasm for integrating physics, biochemistry, and systems biology. J Mol Graph Model 58:1–9.

42. Raschka S, Bemister-Buffington J, Kuhn LA (2016) Detecting the native ligand orientation by interfacial rigidity: SiteInterlock. Proteins Struct Funct Bioinforma 84:1888–1901.

43. Trott O, Olson AJ (2010) AutoDock Vina: improving the speed and accuracy of docking with a new scoring function, efficient optimization, and multithreading. J Comput Chem 31:455–461.

44. Neudert G, Klebe G (2011) DSX: a knowledge-based scoring function for the assessment of protein-ligand complexes. J Chem Inf Model 51:2731–2745.

45. Coleman DE, Sprang SR (1999) Structure of Giα1·GppNHp, autoinhibition in a Gα protein-substrate complex. J Biol Chem 274:16669–16672.

